# Loss of PIK3CA allows *in vitro* growth but not *in vivo* progression of KRAS mutant lung adenocarcinoma in a syngeneic orthotopic implantation model

**DOI:** 10.64898/2026.02.02.701385

**Authors:** Abigail L. Booth, Giuseppe Caso, Barbara Rosati, Ya-Ping Jiang, Wei-Xing Zong, Richard Z. Lin, Harold Bien

**Affiliations:** Department of Physiology & Biophysics, Stony Brook University, Stony Brook, NY, USA; Molecular and Cellular Biology Graduate Program, Stony Brook University, Stony Brook, NY, USA; SBU/Northport VAMC Single Cell and Spatial Multiomic Facility, Stony Brook, NY, USA; Department of Chemical Biology, Ernest Mario School of Pharmacy, Rutgers-The State University of New Jersey, Piscataway, NJ, USA; Northport VA Medical Center, Northport, NY, USA

**Keywords:** PIK3CA, STK11, lung cancer, LUAD, KRAS, orthotopic model

## Abstract

Constitutively active KRAS mutations are highly prevalent in lung cancers, but the direct role of its downstream phosphatidylinositol 3-kinase (PI3K) pathway in tumor progression remains unclear. A previous study established the requirement for PIK3CA, the alpha catalytic isoform, in lung tumor development in mouse models with an intact *Trp53* tumor suppressor. In this study, we further investigated the requirement for PIK3CA for tumor growth both *in vitro* and *in vivo*. We first generated a “KPA” cell line by genetically deleting *Pik3ca* from a murine lung adenocarcinoma “KP” cell line harboring oncogenic *Kras*^*G12D*^ and lacking *Trp53*. We found that *Pik3ca* is not required for cell survival and growth *in vitro*, even under anchorage-independent conditions but reduced the growth rate by 20%. We next orthotopically implanted KP and KPA cells into syngeneic mice and found that PIK3CA is absolutely required for tumor progression, even in the absence of *Trp53*. Implantation of KP cells, or a “KPS” cell line lacking the *Stk11* gene, led to rapid tumor growth and death of all host animals. In contrast, mice implanted with KPA cells all survived with no detectable lung tumors. The gene expression profiles from cultured cell lines suggest KPA cells may be vulnerable to oxidative stress. Indeed, we found KPA cells were more sensitive to hydrogen peroxide and diethyl maleate-induced oxidative stress as compared to KP and KPS cells. Together, these results demonstrate that PIK3CA is not required for lung cancer cell growth induced by mutant KRAS *in vitro* but is critically needed for *in vivo* progression and growth.

## 1. Introduction

Lung adenocarcinoma (LUAD) accounts for about 40% of non-small cell lung cancer (NSCLC) cases and often carries oncogenic *KRAS* mutations, including *KRAS*^*G12C*^, *KRAS*^*G12D*^, and *KRAS*^*G12V*^ [1,2]. While the canonical RAS pathway primarily activates the MAPK signaling cascade, the phosphatidylinositol 3-kinase (PI3K) pathway is also activated downstream of RAS signaling [3-6]. This occurs upon active GTP-bound KRAS binding and signaling through direct binding to the catalytic p110 subunit of the PI3K complex. In solid tumors, the predominant catalytic isoform of the PI3K pathway is the alpha isoform encoded by the *PIK3CA* gene, which is required for tumor growth and maintenance [5-7]. Activated PIK3CA produces phosphatidylinositol-3,4,5 triphosphate (PIP_3_) to activate AKT protein kinases and promotes tumorigenesis and growth in many neoplastic diseases, including lung cancers [5,6,8]. We have previously demonstrated that *Pik3ca* ablation from pancreatic tumor cells results in the clearance of tumor cells by tumor-infiltrating T-cells [3] and similarly, genetic deletion of the downstream *Akt* isoforms dramatically slows tumor growth in murine pancreatic adenocarcinoma [4], further exemplifying the importance of this pathway in cancer biology.

PI3K has been demonstrated to be essential for tumorigenesis and maintenance in a mouse *Kras*^*G12D*^ LUAD model. More specifically, the induced disruption of the binding domain between *KRAS* and *PIK3CA* results in an inability for tumors to grow and causes regression of existing lung tumors [6,8,9]. The PI3K/AKT pathway has been extensively studied as a potential therapeutic target in lung cancer with limited success, partly due to systemic adverse effects of PI3K inhibition [7,10]. Inhibition of the downstream effectors, AKT and mTOR, has been increasingly investigated in recent years as monotherapies, or in combination with PI3K inhibitors, with promising preclinical data [11]. However, clinical trial outcomes in NSCLC patients have been disappointing [12-14] although these trials have not prioritized the selection of biomarkers, such as specific KRAS mutations, which may improve efficacy in specific patient sub-populations. Therefore, more work needs to be done to understand the mechanistic role of *PIK3CA* in tumor progression downstream from *KRAS* binding to define the most appropriate treatment strategies.

Despite the importance of the PI3K/AKT pathway in LUAD, activating mutations in *PIK3CA* are uncommon, especially in *KRAS* mutant LUAD [7,15,16]. In contrast, loss of serine threonine kinase 11 (STK11), also known as liver kinase B1 (LKB1), is frequently found together with oncogenic KRAS mutations in LUAD [17] and typically confers a worse prognosis [1,18]. STK11 is well known to regulate cellular metabolism by activating AMPK in response to hypoxia, nutrient depletion, or increased oxidative stress [18]. AMPK, in turn, inhibits mTOR [18], another downstream effector of the PI3K/AKT pathway, which regulates cell metabolism and growth [19,20]. Here, we constructed and characterized a *Kras*^*G12D*^ *Trp53*^-/-^ mouse LUAD orthotopic implantation model in an immunologically intact host mouse with loss of tumor *Pik3ca* or *Stk11* to explore the effects of each gene in tumor growth and progression.

## 2. Methods and Materials

### 2.1 Cell lines – source and generation

*Generation of KP cells*. The parental mouse lung cancer cell line was derived from tumor-bearing lungs of a *Kras*^*LSL-G12D/+*^*/Trp53*^*flox/flox*^ mouse in C57BL/6J (B6) (Jackson Laboratory) that was infected intranasally with adenovirus expressing Cre recombinase, resulting in expression of one *Kras*^*G12D*^ allele and ablation of both *Trp53* gene copies in lung cells [21,22]. The cells were then infected with a lentivirus expressing firefly luciferase (Cellomics PLV-10064), and clones were selected using G418. The clone with the highest luciferase expression, which we refer to as KP, was selected for subsequent experiments. Tumorigenicity of the KP cells was confirmed by injecting the cells through the chest wall into either lung of syngeneic B6 mice (Supplemental Fig. 1).

*Generation of KPA cells*. KP cells were transfected with CRISPR/Cas9 knockout and Homology Directed Repair (HDR) plasmids (Santa Cruz Biotechnology, sc-422231-KO-2 and sc-422231-HDR-2) to delete the *Pik3ca* gene. Clones were selected in puromycin and loss of PIK3CA expression was verified by western blotting.

*Subcloning of KPA cells*. Cells were reinfected with a lentivirus expressing firefly luciferase (Kerafast FCT230) and clones were selected using blasticidin. All experiments were performed using these selected subclones unless noted otherwise.

*Generation of KPS cells*. KP cells were similarly transfected with CRISPR/Cas9 knockout and HDR plasmids (Santa Cruz Biotechnology, sc-423192 and sc-423192-HDR) to delete the *Stk11* gene. Transfected cells were selected in puromycin and loss of STK11 expression was verified by Western blotting.

*Cell culture*. All cell lines were cultured in RPMI 1640 medium (Corning 10-040-CV) containing 1% penicillin/streptomycin (Gibco 15140-122) and 10% fetal calf serum (Corning 35-015-CV) at 37°C in a humidified atmosphere with 5% CO_2_. Cells were kept in culture for no longer than one month.

### 2.2 In vitro studies

#### Cell proliferation

For assessment of cell growth rates, cells were seeded in cell culture dishes (Falcon #353001) at an initial density of 1.6×10^5^ (Day 0) then counted at 24-hour intervals over 4 days. Growth medium was aspirated and replaced after 48 hours to maintain optimal growth conditions. At each time point, cells were trypsinized (Corning 25-053-CI), an aliquot mixed with an equal volume of trypan blue (Invitrogen T10282) and at least 400 cells were counted with a hemacytometer. Malthusian growth curves were generated and plotted using GraphPad Prism (version 10.5.0).

#### Western blotting

Cells were rinsed twice with cold PBS (Corning 21-031-CV) and then lysed on ice with RIPA buffer containing 50 mM HEPES pH 7.5, 10 mM sodium pyrophosphate, 50 mM NaCl, 50 mM NaF, 5 mM EDTA, 1 mM sodium orthovanadate, 0.25% sodium deoxycholate, 1% NP-40, 1 mM phenylmethylsulfonyl fluoride, and 1 μl/ml protease inhibitor cocktail (Sigma P8340). Cell lysates were then cleared by centrifugation at 14,000 x g for 20 minutes at 4 °C. The protein content of the resultant supernatant was measured using a Bradford assay (BIORAD) on a SpectraMax M5 instrument (Molecular Devices). Equal amounts of protein were combined with SDS sample buffer and separated by SDS-PAGE followed by semi-dry transfer onto nitrocellulose membranes. After blocking in 5% non-fat dry milk, membranes were incubated overnight at 4 °C with primary antibody in Tris-buffered saline plus 0.1% Tween 20. The blot was incubated for 1 hour with Horseradish Peroxidase (HRP)-linked secondary antibody (GE Healthcare) and signals were captured by a FluorChem M imager equipped with AlphaView 3.4.0.0 software (ProteinSimple) using Western Lightning Plus ECL reagents (PerkinElmer NEL105001EA) or SuperSignal West Femto (Thermo Scientific 34095). To detect multiple proteins on one membrane, blots were sometimes cut into pieces that were probed with separate antibodies.

Antibodies to AKT1 (#2938), PIK3CA (#4249), phospho-Ser473 AKT (#4060), AMPK (#2532), and phospho-AMPK (#2531) were purchased from Cell Signaling Technology. Antibodies to STK11/LKB1 (sc-32245) and HSP90 (sc-13119) were purchased from Santa-Cruz Biotechnology.

#### 3D cell culture

Cells were 3D cultured as described by our lab previously [3,4]. Briefly, 1×10^3^ cells were suspended in culture medium with a methylcellulose solution (Sigma M0512-100G) with a final concentration of 0.24%. The cells were cultured by hanging drop method on the lid of a plate filled with PBS to avoid evaporation. The plate was incubated at 37°C with 5% carbon dioxide for seven days and imaged at 40x magnification with light microscopy.

#### MTT assays

Cells were plated at a density of 5×10^3^ cells per well in a 96-well tissue culture-treated flat-bottom plate (Sarstedt 83.3924) and incubated for 48 hours. The cells were then treated with varying concentrations of hydrogen peroxide (Fisher Chemical H325-500), diethyl maleate (DEM) (Fisher AC114440050), cisplatin (Adooq Bioscience A10221), or docetaxel (MedChem Express HY-B0011) for 24 or 48 hours. Cells were also pre-treated for 24 hours with 400 nM alpelisib (MedChem Express HY-15244) or vehicle then the media replaced with 250 µM DEM or 100 µM hydrogen peroxide for an additional 24 hours. 3-[4,5-dimethylthiazol-2-yl]-2,5-diphenyl tetrazolium bromide (MTT) based assays were then performed as previously described [23]. After addition of a 5 mg/mL MTT solution (Alfa Aesar L11939.03), the plates were incubated for 1 hour at 37°C, 5% CO_2_. Formed insoluble formazan crystals were dissolved in DMSO (Thermo Scientific 036480.AP) and the absorbance was read at 570 nm with a 690 nm background subtraction on a Spectromax M5 (Molecular Devices).

#### Statistics

All statistics were calculated using GraphPad Prism using a one or two-way ANOVA with a Bonferroni post-hoc test and a significance value of 0.05. Survival curves were analyzed in the same software with a log-rank test.

### 2.3 *In vivo* studies

#### Orthotopic implantation of tumor cells in the lung

All procedures involving live animals were performed according to institutional policies at Northport VAMC (ACORP) and Stony Brook University (IACUC).

Cells to be used for orthotopic implantation were trypsinized and extensively washed with cold PBS by repeated centrifugations at 300 x g for 5 minutes at 4 °C. The number of viable cells was then counted in the presence of Trypan Blue using a Countess II Automated Cell Counter (Invitrogen). The injection solution was prepared by suspending 1×10^7^ viable cells per mL in PBS containing 0.5 mg/mL of Matrigel (Corning 356231) and kept on ice until implantation. All steps were carried out under aseptic conditions.

Cells were implanted in the lung of 8-to 14-week-old male C57BL/6J (B6) mice or SCID (B6.CB17-Prkdc^scid^/SzJ) mice purchased from Jackson Laboratories. Mice were anesthetized with 2-4% isoflurane. A 5 mm incision was made on the skin over the left lung, and the underlying muscles were separated to visualize the ribs and motion of the left lung unless otherwise specified. A 28-gauge needle attached to a ½ cc U-100 insulin syringe was inserted to a depth of about 3 mm at the mid-axillary line and 50 μl of injection solution containing 5×10^5^ cells was injected into the upper left lung. The wound was closed with a 9 mm wound clip. Post-procedure, mice were injected subcutaneously with 2 mg/kg ketolorac for pain relief and 0.1 ml of 2.27 mg/ml enrofloxacin antibiotic. After implantation, the animals were observed for 30-60 minutes to monitor their recovery.

#### Animal Survival Studies

After implantation, mice were monitored for signs of distress and weighed at least weekly or more frequently if persistent weight loss was noted. Mice were euthanized if they lost 15% or more of their initial weight and/or appeared to be in severe distress as determined by a blinded observer. A necropsy was performed to confirm the presence of gross tumors. For some experiments, tumors or lungs were subsequently dissected, fixed for 24 hours in a 10% buffered formalin solution, and then embedded in paraffin blocks. Kaplan-Meier survival curves were generated and plotted using GraphPad Prism.

#### In-vivo luminescence imaging (IVIS)

For the initial tumor growth and progression study of the parental KP cells, tumor progression was monitored using a Lumina III In Vivo Imaging System (IVIS, PerkinElmer) as previously described by us [3].

#### H&E staining

H&E staining of suspected tumors fixed and embedded in paraffin blocks was performed at HistoWiz, Inc, using the Leica Bond RX automated stainer (Leica Microsystems) and a standard protocol. Blocks were sectioned at 4 μm then slides were dewaxed using xylene and alcohol based dewaxing solutions. Tissue was incubated with hematoxylin according to the manufacturer’s protocol. The slides were visualized using an Aperio GT 450 DX slide scanner (Leica Biosystems) at 40X. Resultant images were analyzed for identification of tumor sections.

### 2.4 Bulk-RNA sequencing

Cells were cultured to 80-90% confluence in 10 cm cell culture dishes (Sarstedt 83.3902), washed with cold PBS, and lysed using 600 µL RLT Plus buffer (Qiagen 1030963) +1% v/v ß-mercaptoethanol (Sigma Aldrich M6250-100ML) and the lysate homogenized using QIAshredder (Qiagen 79654), following the manufacturer’s instructions. Total RNA was extracted using the Qiagen AllPrep DNA/RNA/miRNA Universal Kit (Qiagen 80224) and quantified using a Nanodrop 2000 (ThermoFisher).

RNA integrity was analyzed on a TapeStation 4200, and the RNA Integrity Numbers (RINs) were 9.8 and higher for all samples used. Bulk RNA library preparation and sequencing was performed by a commercial supplier (Novogene, Inc.), using the NEBNext® Ultra™ II RNA Library Prep Kit for Illumina (New England Biolabs E7770) with the NEBNext® Poly(A) mRNA Magnetic Isolation Module (New England Biolabs E7490). Libraries were sequenced at a depth of approximately 33 million paired reads per sample (PE150) on an Illumina NovaSeq X Plus Instrument.

#### RNA-seq Data Analysis

The transcripts were quantified and mapped to the GRCm39 (GCA_000001635.9) reference genome from ENSEMBL using Salmon with correction for GC bias. The resultant quantification files were loaded into R (version 4.5.1) in RStudio (version 2024.12.1, build 563) and the DESeq2 package (version 1.48.1) [24] identified differentially expressed genes. To produce a heatmap with the most differentially expressed genes between cell types with the least variation between replicates with the geneFilter package (version 1.90.0), the data was transformed with a variance-stabilizing transformation, but all further analyses were done with the untransformed DESeq2 results to limit bias. The results were filtered to only include genes with a *p*-value less than 0.01, an absolute log_2_-fold change greater than one, and a count of at least 250. This filtered data was used with the gseGO function of clusterProfiler (version 4.16.0) [25] and fgsea (version 1.34.2) to find the gene ontology enriched by molecular function with a more relaxed *p*-value limit less than 0.05.

## 3. Results

### 3.1 Loss of Pik3ca, but not Stk11, moderately reduces in vitro growth rate in murine Kras mutant lung adenocarcinoma cell line

To study the effect of loss of PIK3CA in mutant KRAS lung cancer, we developed a murine tumor cell line (KPA) lacking this protein from a parental cell line (KP), which we obtained through isolation of lung tumor cells from a *Kras*^LSL-G12D/+^/*Trp53*^flox/flox^ B6 mouse after intra-nasal infection with Cre expressing adenovirus and subsequent transfection with a luciferase gene to non-invasively track longitudinal tumor progression. A CRISPR-Cas9 system was then used to genetically delete *Pik3ca* from KP cells to create the “KPA” cell line. We also used the same CRISPR-Cas9 technique to produce a *Stk11*-deficient “KPS” cell line from KP cells to study the effects of loss of STK11 in mutant KRAS lung cancer. We validated PIK3CA and STK11 deletion by western blotting (Fig. 1A). Additionally, deletion of the target genes resulted in decreased phosphorylation of AKT in KPA or AMPK in KPS cell lines, respectively (Fig. 1A), consistent with canonical signal transduction from PI3K to AKT [3,7,9,26] and STK11 as one of the kinases that phosphorylates AMPK [18].

**Figure 1.**
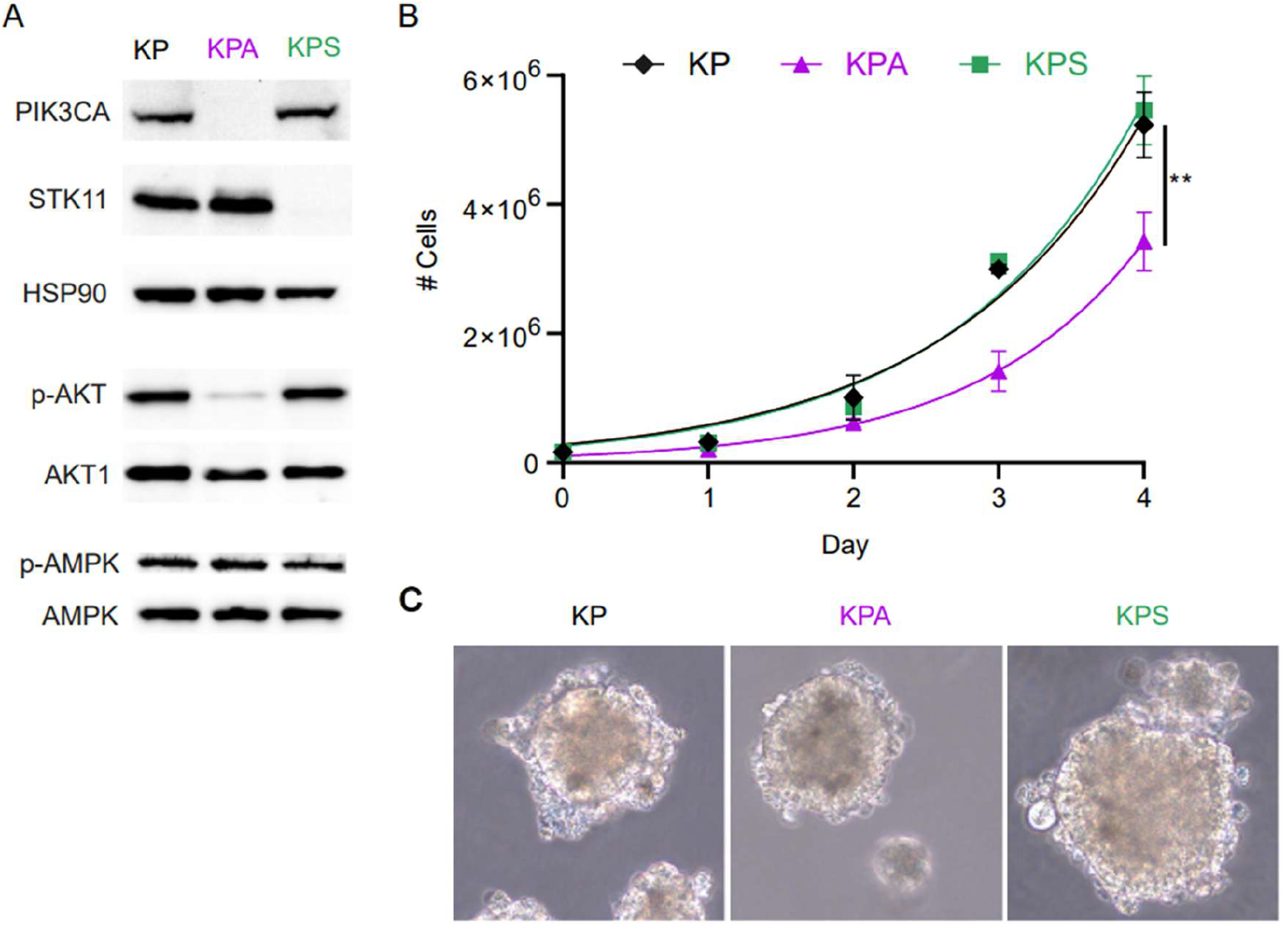
Generation and in-vitro characterization of KP, KPA, and KPS cell lines. (A) Western blot of parental KP, derived KPA (Pik3ca^-/-^), and KPS (Stk11^-/-^) cells with HSP90 as loading control. (B) Proliferation of KP, KPA, and KPS cells *in vitro*. Shown are averages of 3 separate experiments with the standard error of the mean. Growth at each day compared by two-way ANOVA with Bonferroni post-hoc for day 4 (KP vs KPA p = 0.01). (C) Representative light-microscopy images of spheroid growth of KP, KPA, and KPS cell lines after 7 days of growth in 3D-methylcellulose culture (40X magnification).

We first measured the *in vitro* proliferation rates over several days and compared them to the growth of the parental cell line, KP (Fig. 1B). While the KP and KPS cells maintained a nearly identical doubling time of 0.94 and 0.90 days respectively (95% confidence interval (CI) 0.80-1.10 and 0.77-1.1), the growth rate of the KPA cells diverged at day 2 with an overall 20% slower doubling time of 0.79 days (95% CI 0.67-0.94) at day four when compared to the KP and KPS cell lines (Fig. 1B). Next, we tested the ability of each cell line for anchorage-independent growth by growing the three cell lines in a methylcellulose-based 3D culture, as previously described [3]. Over the course of a week, all the cell lines grew into spheroids, indicating that while the cells are adherent in culture conditions, anchorage is not a requirement for cell growth in any of the cell lines (Fig. 1C).

### 3.2 Loss of PIK3CA but not STK11 markedly decreases tumor growth and progression in vivo

To investigate the distinct roles of *Pik3ca* and *Stk11* in a physiologically relevant model of LUAD, we implanted the three cell lines orthotopically into the lungs of mice. Identical numbers of KP, KPA, or KPS cells were implanted into syngeneic B6 mice in several independent experiments and monitored without further treatment until death or measurable, pre-specified end points, including excessive weight loss or signs of distress.

KP and KPS injected mice had similar overall survival (Fig. 2A) with KP injected mice surviving a median of 25.5 days (95% CI 20-28 days) and KPS injected mice surviving a median of 28 days (95% CI 25-37 days). In contrast, none of the KPA injected mice died or reached pre-specified endpoints before the end of the study (82 days post-implantation) (Fig. 2A).

**Figure 2.**
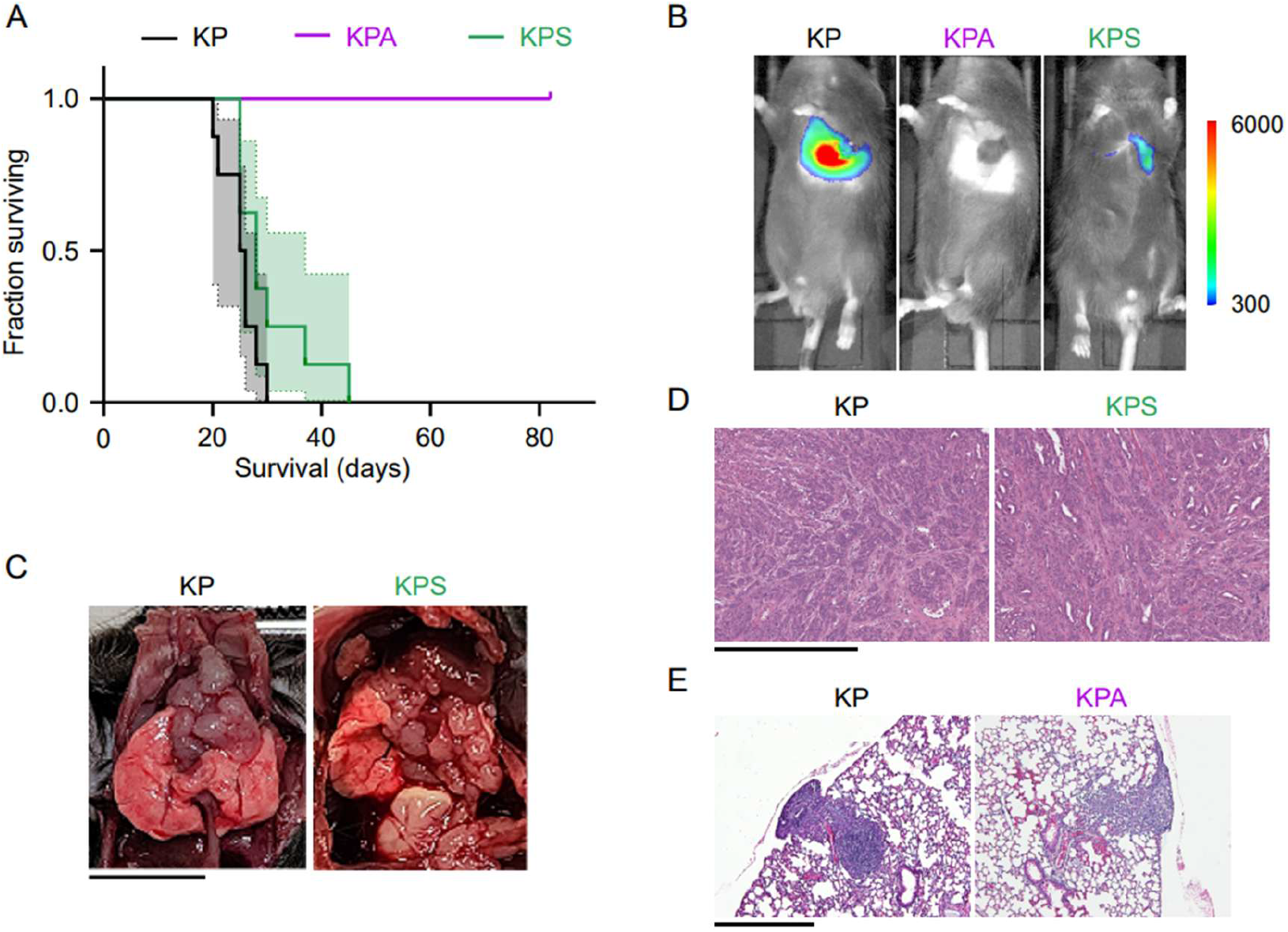
*In vivo* tumor progression of implanted KP, KPA, and KPS cells in murine lung adenocarcinoma model. 5×10^5^ KP, KPA, or KPS cells were implanted into the lungs of mice in several independent experiments. (A) Kaplan-Meier survival curves for mice implanted with KP (n = 8), KPA (n = 7), or KPS (n = 8). Median survival: KP, 25.5 days; KPS, 28 days; all KPA alive on day 82. p < 0.0001, KP versus KPA (log-rank test). (B) Representative tumor progression observed with IVIS imaging 18 days post-implantation. Scale represents luminescence intensity (arbitrary units). (C) Representative post-mortem analysis of sacrificed KP and KPS implanted mice. Scale bar represents 10 mm. (D) Representative H&E staining confirming tumors in KP and KPS-implanted mice. Scale bar represents 500 µm. (E) To assess early tumor development, 5×10^5^ KP or KPA cells of earlier subclones were implanted into the lungs of syngeneic mice (n = 3 mice per group). At 10 days post-implantation, the lungs were fixed and sectioned. Shown are representative H&E stained sections of lungs from each group of mice. Scale bar represents 500 µm.

*In vivo* fluorescence imaging was used to detect the luciferase signal from implanted cells for non-invasive, weekly tumor monitoring post-implantation with In-Vivo Imaging System (IVIS) imaging (Fig. 2B). At 18 days post-implantation, KP and KPS injected mice typically exhibit obvious tumor development localized around the lungs, albeit with a stronger luminescent signal in mice injected with KP cells than KPS cells (Fig. 2B). Sacrificed mice that exhibit a strong luminescent signal on IVIS reveal extensive development of large masses around the lungs, mediastinum, and diaphragm (Fig. 2C) which can be confirmed through microscopic examination to be tumors with similar morphology between KP and KPS implanted mice (Fig. 2D). Conversely, KPA injected mice show no clear evidence of macroscopic tumor growth through the end of the study (Fig. 2B).

Since the KPA cells were previously demonstrated to have the ability to grow anchorage-independently in culture, we looked more specifically at the early lung tumor development for the KPA cells to confirm that the tumors could engraft in the lung. To do this, we harvested the lungs of mice orthotopically implanted with KP or KPA cells at 10 days post-implantation (Fig. 2E). None of the KP or KPA-injected mice had visible tumors on the surface of the lungs (not shown), but microscopic evaluation of the H&E-stained sections revealed clear tumor growth in all mice implanted with KP cells and in 2 out of 3 mice implanted with KPA cells (Fig. 2E).

This result resembles our previous findings, where PIK3CA ablation resulted in T cell-mediated clearance of pancreatic tumors, and therefore, improved survival [3]. To investigate if a similar immune mechanism is occurring in this lung cancer model, we implanted KPA cells into SCID mice in the B6 genetic background that lack a functional adaptive immune system. SCID mice implanted with KPA cells also did not form tumors (not shown), and all host animals survived until the end of the study, like the KPA-implanted B6 mice (Supplemental Fig. 2A). To rule out that these results were due to a requirement for a higher starting number of implanted KPA cells, we repeated the experiment with a 10-fold higher cell count and found the equivalent 100% host animal survival (Supplemental Fig. 2B). This indicates that, although KPA tumors can engraft in the lungs of mice (Fig. 2E), they fail to progress by some mechanism distinct from adaptive immune clearance. In summary, these results align with the previously established requirement of PIK3CA binding to KRAS for *in vivo* tumor progression [6,8] but raise additional questions on the mechanism for the KPA cells’ ability to grow *in vitro* but not progress *in vivo*.

### 3.3 Cells deficient in Pik3ca have reduced gene expression in key oxidative stress pathways

To identify potential mechanisms underlying the lack of *in vivo* lung tumor progression of KPA tumor cells, we compared the gene expression profiles of cultured KP, KPA, and KPS cells using bulk RNA sequencing. We transformed the data with a variance stabilizing transformation to identify the most differentially expressed genes (DEG) between cell types with the least variation between replicates to yield the most robust, consistent changes. Next, we performed gene set enrichment analysis (GSEA) to determine if there were any other molecular changes that were significantly different between the cell lines. When all DEGs were included in this analysis, the most changed pathways in either KPA or KPS in comparison to KP were identical (data not shown), so we stringently filtered the gene list to prioritize DEGs with a large, highly significant log_2_-fold change compared to KP cells to ensure we only included robust and specifically different genes in GSEA.

From the remaining list of 1830 and 592 DEGs in KPA or KPS cells, respectively, we then analyzed the enriched molecular function gene ontology terms to identify differences between molecular processes which may contribute to the disruption of *in vivo* tumor growth. KPA cells had decreased activity in both peroxidase and antioxidant pathways (Fig. 3A) while KPS cells only had two significantly activated functions of monosaccharide and carbohydrate binding (Fig. 3B). This suggests that loss of *PIK3CA*, but not *STK11*, from LUAD tumor cells may affect the redox balance and/or the metabolic response to oxidative stress.

**Figure 3.**
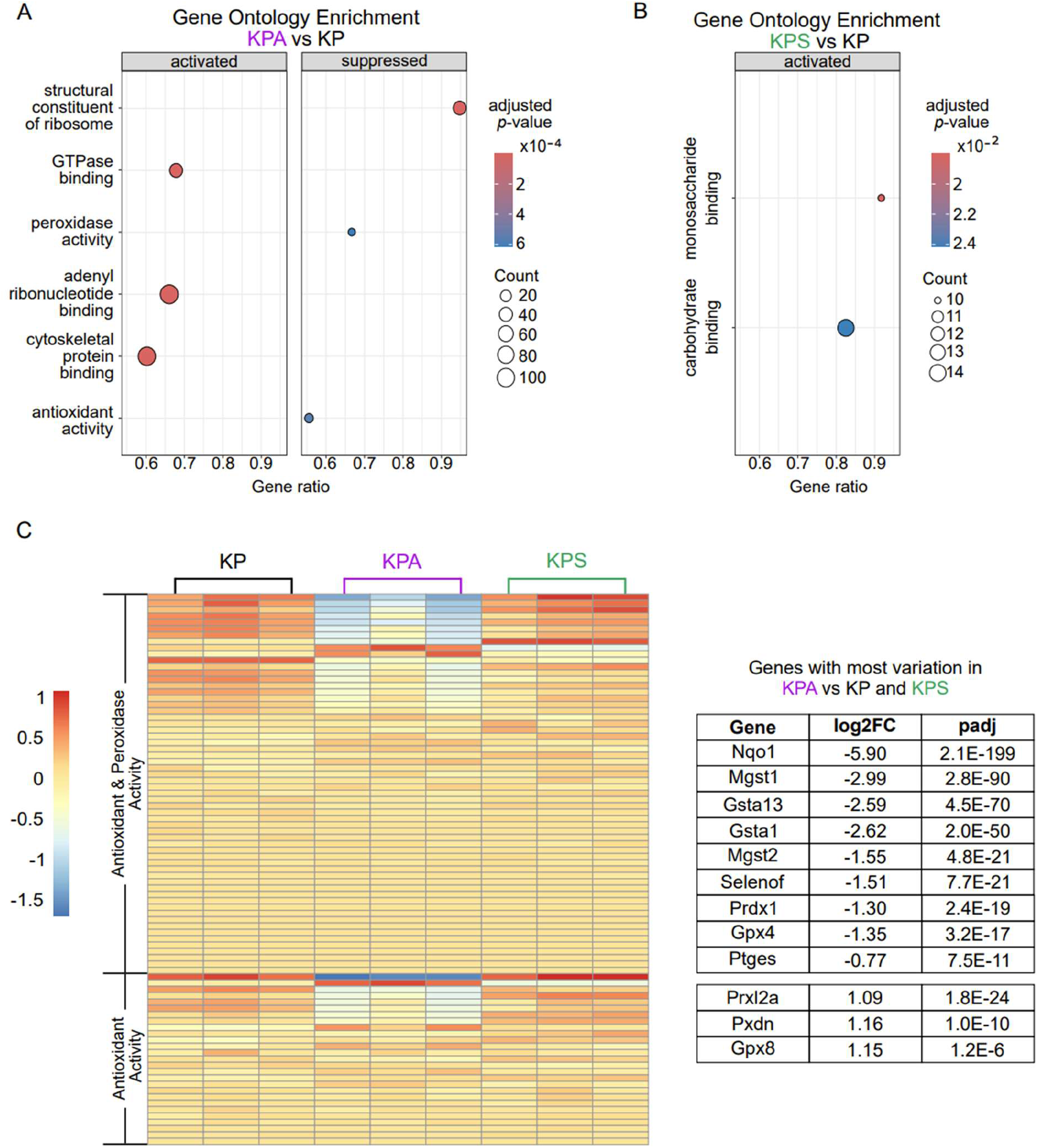
RNA sequencing comparison of pathway enrichment. (A-B) RNA was harvested from cultured cell lines and sequenced. Gene Set Enrichment Analysis (GSEA) analysis of the top activated and/or suppressed molecular functions by gene ontology in KPA (A) and KPS (B) compared to KP. (C) Heatmap of variance-stabilized differential expression of all genes in the antioxidant activity (GO:0016209) and peroxidase activity (GO:0004601) pathways (left). The genes with the greatest variability in expression in the KPA cell line in comparison to both other cell lines are presented in the table on the right with the log_2_ fold change (log2FC) and adjusted *p*-value (padj) in comparison to the KP cell line.

To gain more insight into how KPA cells’ peroxidase and antioxidant gene expression differed from the KP and KPS cell lines, we compared the expression of all genes within these pathways. Peroxidase activity is a subset of antioxidant activity, so only overlapping genes specific to the antioxidant activity pathway are presented. Approximately half of the genes had any significant variation between the three cell lines and, in accordance with the overall downregulation of both pathways seen in Figure 3A, most of these genes are downregulated in KPA cells compared to KP cells (Fig. 3C). For the genes with the greatest variation between all three cell lines, KPA cells had consistently opposite direction of gene expression changes to both KP and KPS cells. This suggests KPA cells may have a reduced ability to manage reactive oxygen species (ROS) and maintain redox balance, which may contribute to the inability of these cells to generate tumors *in vivo* (Fig. 3C). This is supported by the specific decrease in gene expression for several downstream targets of the transcription factor NRF2, a redox-responsive regulator of antioxidant and detoxification responses, including glutathione transferases (*Mgst1, Gsta1, Mgst2, Gsta13*), a lipid hydroperoxide-reducing glutathione peroxidase (*Gpx4*), and oxidoreductases (*Nqo1* and *Prdx1*) [27,28] (Fig. 3C).

### 3.4 Inhibition of PIK3CA activity increases sensitivity to oxidative stress but enhances resistance to cytotoxic chemotherapies

ROS have a well-established involvement in KRAS-driven lung cancers, and oncogenic Ras, including KRAS, is well known to promote constitutive production of ROS [29]. In addition, lung parenchyma has a higher oxygen concentration than any other organ and thus contains additional regulatory mechanisms for oxygen sensitivity [30,31]. Therefore, lungs are proposed to have specialized responses to ROS in comparison to other organs, although the details of this are incompletely understood [31,32]. Based on the mRNA expression changes previously discussed, we hypothesized that the KPA cells could be more sensitive to oxidative stress due to a reduced ability for ROS detoxification.

To test the sensitivity of each cell line to ROS, we treated KP, KPA, and KPS cells *in vitro* with hydrogen peroxide (H_2_O_2_). Extracellular treatment with H_2_O_2_ increases intracellular ROS levels and, at high doses, induces cell death [33]. Therefore, we exposed KP, KPA, and KPS cells to varying concentrations of H_2_O_2_ and measured the residual metabolic activity using MTT assays. While KPA cells were markedly more sensitive to H_2_O_2_ than KP cells, KPS cells were dramatically less sensitive (Fig. 4A). For KP cells, loss of approximately 50% metabolic activity (IC_50_) compared to control-treated KP cells occurred around 82.5 µM of H_2_O_2_ (95% CI: 66.1 to 104). In KPA cells, IC_50_ was approximately 35.1 µM of H_2_O_2_ (95% CI: 28.7 to 42.7), meaning that KPA cells are about 43% more sensitive to ROS than KP cells. At the highest tested concentration of 100 µM H_2_O_2_, KPS cells only decreased to 76% metabolic activity with respect to control-treated KPS cells, making the IC_50_ indeterminable from this data (Fig. 4A). This aligns with previous studies demonstrating that *STK11*-deficient mouse embryonic fibroblasts increase intracellular ROS accumulation levels by two-fold compared to wild-type cells in response to treatment with a sub-cytotoxic dose of H_2_O_2_ [34].

**Figure 4.**
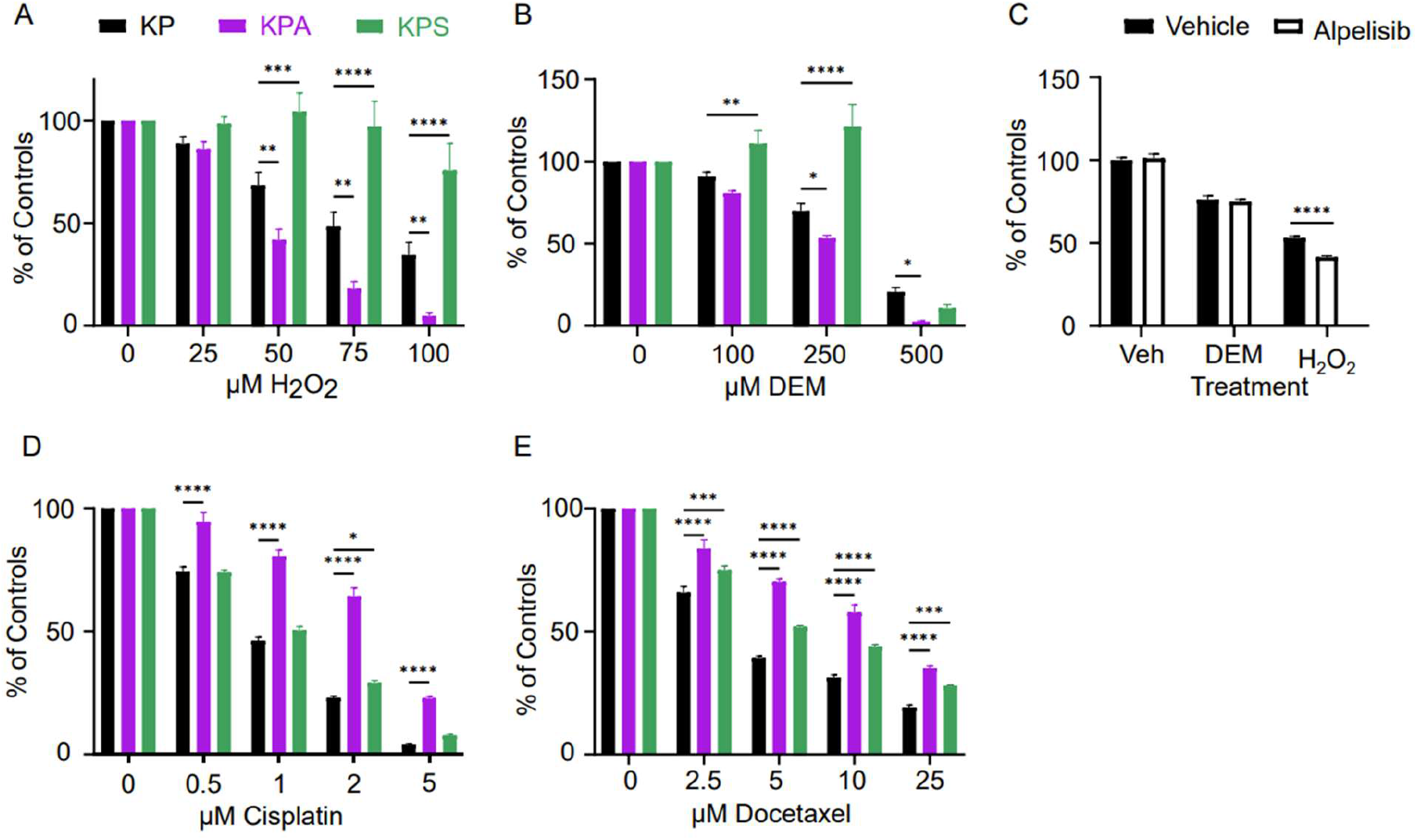
Effects of hydrogen peroxide, chemotherapy, and glutathione depletion on KP, KPA, and KPS cells. (A-B) Cell lines were treated with a dose-response of hydrogen peroxide (A) or diethyl maleate (DEM) (B) for 24 hours and the metabolic activity of the cells was measured by MTT assay. Plotted is the mean and standard error of absorbance at 570 nm from five replicates across 4 independent experiments. (C) KP cells were pre-treated with 400 nM alpelisib or vehicle for 24 hours then treated with 250 µM DEM, 100 µM hydrogen peroxide, or vehicle for an additional 24 hours and the metabolic activity measured with MTT assay. Plotted is the mean and standard error of absorbance at 570 nm from twenty replicates. (D-E) Cell lines were treated with a dose-response of cisplatin (D) or docetaxel (E) for 48 hours and the metabolic activity assessed by MTT assay. Plotted is the mean and standard error of absorbance at 570 nm from 5 replicates across 4 independent experiments. In all experiments, the variance at each dose was compared against KP with a two-way ANOVA and Bonferroni post-hoc * p < 0.05; ** p < 0.01, *** p< 0.005, **** p < 0.0001.

Glutathione acts as a reducing agent to detoxify ROS and maintain redox balance within the cellular environment [35]. Diethyl maleate (DEM) is a commercially available electrophile that alkylates reduced intracellular glutathione to block its antioxidant activity [33,36,37]. Therefore, treatment with DEM can inhibit cellular antioxidant response, and hence, potentially increase sensitivity to ROS. Our results revealed that both KP and KPA cells had a dose-dependent response to DEM treatment with the IC_50_ for KP being 353 µM (95% CI: 290 to 423) and KPA being significantly more sensitive with an IC_50_ of 201 µM (95% CI: 169 to 237) (Fig. 4B). While the KPS cells also showed some dose-dependent response to DEM, the behavior was very different with metabolic activity slightly increasing with DEM treatments up to a concentration of 250 µM and then plummeted to 10.7% metabolic activity of the control with the 500 µM dose, possibly due to a shift from oxidative to glycolytic metabolism to account for decreased mitochondrial spare respiratory capacity [38] (Fig. 4B).

To determine if the lack of PI3K catalytic activity is sufficient to explain the altered response to ROS in KPA cells, we pharmacologically inhibited PIK3CA activity in KP cells with alpelisib. This drug, commercially known as Piqray®, is a potent selective PIK3CA inhibitor, which blocks the kinase activity of PIK3CA and is currently used in the treatment of breast cancer [39,40]. We treated KP cells with DEM and H_2_O_2_, with and without alpelisib, to test whether inhibition of PIK3CA kinase activity would be sufficient to promote ROS sensitivity of KP cells— either directly or via glutathione depletion— to the levels seen in KPA cells. At a dose of 250 µM DEM, there is no difference in the metabolic activity levels of KP cells with or without alpelisib treatment (Fig. 4C). By contrast, alpelisib treatment caused the KP cells to be significantly more sensitive to 100 µM H_2_O_2_ than vehicle-treated control cells (Fig. 4C). The combined treatment of alpelisib and H_2_O_2_ only resulted in a 58% loss in metabolic activity (Fig. 4C) whereas treatment of KPA cells with the same concentration of H_2_O_2_ yielded a 95% loss (Fig. 4A). Taken together, these findings suggest that pharmacological inhibition of PI3K kinase activity alone is insufficient to completely describe the effect of loss of PIK3CA expression in ROS sensitivity.

We next tested whether the increased sensitivity to drug treatments in KPA cells was a generalized, nonspecific drug effect by testing sensitivity to traditional cytotoxic chemotherapeutic agents. We treated KP, KPA, and KPS cells with different concentrations of cisplatin, a key component of cytotoxic chemotherapy for NSCLC [41-44]. Treatment of each cell line with cisplatin demonstrated strong resistance of KPA cells to the drug in comparison to KP and KPS cells, which demonstrated similar levels of sensitivity to each other (Fig. 4D). The estimated concentration of cisplatin which reduced metabolic activity to 50% of control untreated KP cells (IC_50_) for KP and KPS cells overlapped with IC_50_ values of 0.85 (95% CI: 0.72 to 1.0) and 0.98 µM of cisplatin (95% CI: 0.86 to 1.1), respectively (Fig. 4D). In contrast, KPA cells were significantly more resistant to cisplatin with an IC_50_ of 3.0 µM (95% CI: 2.4 to 3.7).

Similarly, we also exposed cells to docetaxel, a member of the taxane family and another commonly employed chemotherapy for NSCLC [45]. Treatment of KP, KPA, and KPS cells with docetaxel resulted in a dose-dependent decrease in metabolic activity across all cell lines (Fig. 4E). For KP cells, the IC_50_ of docetaxel was 4.3 µM (95% CI: 3.8 to 4.7). KPS cells had a slightly higher IC_50_ of 7.1 µM (95% CI: 6.4 to 7.8), indicating slightly less sensitivity to docetaxel compared to KP at all tested concentrations. Like cisplatin, KPA cells showed significant resistance to docetaxel with an IC_50_ of 13 µM (95% CI: 12 to 15), over 300% less sensitive to docetaxel than KP cells (Fig. 4E). In summary, KPA cells exhibited increased sensitivity to ROS stress but were more resistant to traditional cytotoxic chemotherapy compared to KP or KPS cells. This suggests that increased sensitivity to ROS from loss of PIK3CA activity is not a general increase in sensitivity to cellular stress but rather a specific response to ROS.

## 4. Discussion

The importance of the PI3K pathway in many cancers, including lung [7], pancreatic [3,46], colorectal [16], and cervical cancers [47], is well known. Previous efforts in understanding PIK3CA in KRAS-driven NSCLC have largely focused on the clinically relevant overactive *PIK3CA* phenotype from well characterized mutations [7,15,16,39,48], but disruption of PIK3CA signaling in a mouse model has also revealed its necessity for tumor growth and maintenance [6,8,9]. More specifically, disruption of the RAS-binding domain of PIK3CA abrogates tumor progression and reduces the size of existing lung tumors [6]. This suggests that pharmacological inhibition of PIK3CA might be a viable treatment option for *KRAS*-driven cancers. However, clinical attempts at pharmacologically inhibiting PI3K activity have been unsuccessful due to toxicity of systemic PI3K inhibition and possible putative compensatory activation of alternative cell signaling pathways [7,10,49] indicating the need for further investigation regarding the broader implications of reduced *PIK3CA* expression and/or activity. Therefore, we established and characterized a murine syngeneic orthotopic implantation *Kras* mutant *Trp53* deficient LUAD model to better understand the role of PIK3CA within the context of LUAD. To apply the model, we developed a KRAS mutant PIK3CA-null KPA tumor cell line. We also looked at STK11, whose loss is well established to result in more aggressive LUAD through shifting metabolic dependencies [1,18], with a KRAS mutant STK11-null KPS tumor cell line.

Previous methods to test the requirement for PI3K activity in lung cancer have involved mouse models with germline deletions or inducible modifications of PIK3CA [6,8,9]. These models were instrumental to the initial understanding of how KRAS and PIK3CA bind and interact to transduce signaling and affect *in vivo* tumor growth but have limited potential for further *in vivo* studies and are unable to examine the mechanisms precisely. Additionally, loss of function mutations in *TP53* are frequently seen in conjunction with *KRAS* mutation in human LUAD and results in 30% worsened overall survival [50]—a finding not mimicked in these models, which also limits the ability to generate cell lines from resultant tumors [51]. *In vitro* studies with primary embryonic fibroblasts from mice with mutated PIK3CA RAS-binding domains report accumulation of cells in G1 and G2 phases of mitosis and reduced cell division in comparison to wild-type cells. The mutated fibroblasts lose the capacity for anchorage-independent growth [8], so our ability to generate and grow the KPA cell line was unexpected and indicates that PIK3CA is not a requirement for *in vitro* growth. Unlike KP and KPS cells with similar growth rates, we observed a small but significant decrease in *in vitro* growth rate of KPA cells compared to KP cells. Furthermore, both KPA and KPS cells demonstrated anchorage-independent growth. This provides an interesting potential opportunity to study the role of PIK3CA in lung cancer, given that similar models of pancreatic [3] and lung cancer with silenced oncogenic *KRAS* [52] failed to grow *in vitro* or *in vivo*.

Therefore, we took advantage of the demonstrated *in vitro* growth ability and orthotopically implanted the KP, KPA, and KPS cells into the lungs of wild-type B6 mice with intact immune systems to investigate the growth rates in a more biologically relevant context. Although KP and KPS tumors grew at similar rates with similar survival curves, KPA tumors failed to progress *in vivo* resulting in a marked increase in survival in comparison to KP and KPS-injected mice. The observed reduced *in vitro* growth rate alone is insufficient to account for the prolonged survival and lack of tumors. Additionally, KPA cells are able to successfully implant into the lungs yet are unable to progress even in the absence of a functional adaptive immune response in contrast to a similar murine model of pancreatic cancer [3,53]. Taken together, this indicates that *PIK3CA*, but not *STK11*, is required for growth and progression *in vivo* in *KRAS* mutant NSCLC but dispensable for *in vitro* growth. This contrasts with activating KRAS mutations where our efforts to revert the oncogenic KRAS mutation present in KP cells back to wild-type were met with failure due to lack of *in vitro* growth (data not shown). Therefore, we began investigating the potential mechanisms driving the observed differences between *in vivo* and *in vitro* behavior of KP, KPA, and KPS tumor cells with a focus on the KPA cells.

By analyzing the transcriptomic profiles of the cultured cell lines, only the KPA cells presented reduced expression across peroxidase and antioxidant activity pathways, indicating a potential vulnerability to redox stress. Redox homeostasis is essential in healthy cells to maintain the balanced production of ROS, generated in the mitochondria as a byproduct of aerobic cellular respiration, with degradation by antioxidants to avoid excessive oxidative stress [33,36,54]. Cancer cells often have higher endogenous ROS levels than non-cancerous cells and use this as a survival mechanism to promote proliferation, but there exists a threshold of ROS at which the benefits are surpassed and cells will die [55]. Accordingly, ROS can have either pro- or anti-tumor effects depending on the local concentration and specific microenvironment of the tumor [56]. Induction of excessive ROS in cancer cells has been reported to reduce cell proliferation in a PI3K/AKT-dependent manner [18,55,57]. Therefore, accumulation of ROS to cytotoxic levels is a mechanism for killing aberrantly proliferative cells and a common indirect therapeutic strategy to halt tumor progression in oncology [54]. However, in many cancers, metabolic reprogramming is utilized by tumor cells to generate additional antioxidants and redox cofactors in the increasingly stressed, hypoxic microenvironment generated through malignant tumorigenesis [33,37,48,54,58-60]. Based on the genes that varied most from the KPA cell line in comparison to the other cell lines, there may be a deficiency in NRF2 sensing of oxidative stress and activation of antioxidant enzyme expression [27]. KRASoncogenic mutations have been demonstrated to promote tumor development through activation of antioxidant genes by NRF2 [61], making this a potential mechanism or the lack of KPA tumor progression *in vivo*. However, many of these genes have diverse functions in the cell, such as *Nqo1*, which acts as a superoxide reductase, but also associates with microtubules during mitotic spindle formation [62]. Additionally, some of the downregulated genes in KPA cells, particularly *Gpx4*, point toward ferroptosis or autophagy inhibition as other potential mechanisms [61,63]. More work is required to clearly identify the precise cellular process that differs in KPA cells from KP and KPS cells resulting in failure to progress *in vivo*.

We found loss of PIK3CA strikingly increased sensitivity to ROS via hydrogen peroxide exposure, in contrast to loss of STK11 which exhibited marked resistance when compared to the parental KP cells. KRAS overactivation, which exists across all tested cell lines, is known to induce tumorigenicity with requirements for mitochondrial production of ROS and glutamine catabolism [64-66]. Similarly, depletion of glutathione, an antioxidant produced from glutamine [67], with DEM resulted in the same inverse effects on metabolic viability between KPA and KPS cells. DEM increases ROS generation through glutathione depletion and inhibits the cell cycle [68]. While little is known about the specific effects of decreased expression of PIK3CA on metabolic changes in cancers, there is existing evidence that hyperactivated PIK3CA shifts glutamine metabolism in cancer [48,60,67]. Across multiple human cancer cell lines, hyperactivated PIK3CA in tumors requires 2-oxoglutarate dehydrogenase for survival and proliferation due to an increased glucose metabolism and reduced NAD+/NADH ratio [48]. Accordingly, glutathione biosynthesis, simulated by oncogenic PI3K/AKT signaling, has been shown to trigger the metabolic reprogramming in breast cancer [67]. By contrast, STK11 has a well-documented role in metabolic reprogramming [18,38,58,59]. STK11 regulation of AMPK is essential for maintaining nicotinamide adenine diphosphate (NADPH) homeostasis and increasing glucose catabolism [59], whereas loss of STK11 increases HIF-1α with increased mTOR expression and higher ROS, which results in a more glycolytic-dependent metabolism [58]. In support of our findings, *KRAS* mutation with *STK11* deletion has been reported to maximize metabolic capacity, causing cells to be more resistant to oxidative stress [38].

Interestingly, pharmacological inhibition of PIK3CA kinase activity with alpelisib sensitized KP cells to ROS but not to the same degree as complete ablation of the protein. Additionally, the drug did not affect the metabolic response to glutathione depletion. This suggests that there may exist some additional mechanism affecting metabolic signaling in KPA cells beyond the kinase activity of PIK3CA, including possibly a structural role of PIK3CA itself. Therefore, studies that disrupt the kinase activity [9] of PIK3CA or even loss of the RAS-binding domain in PIK3CA [6] may not be sufficient to elucidate its role in tumor development. Future studies are required to further investigate potential non-kinase and non-KRAS binding domain roles for PIK3CA in tumor cell growth *in vitro* and *in vivo*.

Furthermore, KPA and KPS cells both demonstrated significant resistance to first-line NSCLC chemotherapies which aligns with the clinically established chemotherapy-resistant nature of KRAS mutant NSCLC with loss of STK11 function [69]. The increased resistance of KPA suggests that complete inhibition of the PI3K pathway in KRAS mutant NSCLC with wild-type PIK3CA may induce resistance to traditional cytotoxic chemotherapy, possibly due to decreased mitotic rates. However, the lack of *in vivo* growth of KPA cells with loss of PIK3CA suggests a contradictory effect. Future studies using KPA cells could investigate these two distinct facets of PIK3CA in KRAS mutant NSCLC. Several orthotopic implantation models for NSCLC have previously been described [70,71]. However, these have involved the intrathoracic implantation of a xenograft [70] or used intratracheal delivery of organoids or cells, which requires bleomycin pre-treatment for successful cell delivery [71]. Therefore, our model benefits from the use of syngeneic cells directly implanted into the lung without anti-mitotic chemotherapy treatment for the least disruption to the host lung microenvironment. Through the implementation and characterization of this model, we have demonstrated its utility with the implantation of our cell line lacking PIK3CA having the novel ability to grow *in vitro* but not *in vivo*, while our KP and KPS cell lines exhibited similar *in vitro* and *in vivo* growth. Our preliminary studies suggest a sensitivity of the KPA cell line to ROS in comparison to parental KP or KPS cells that may affect tumor development in the lung but not encountered in standard cell culture conditions. One potential difference in the *in vitro* environment could relate to the aerobic composition of the microenvironment in a lung. However, the ratio of partial pressure of oxygen between a tumor and its surrounding lung tissue has been measured to have high variability between human NSCLC patients [72], making this a potentially complicated feature to mimic. Some cell culture strategies, including air-liquid interface [73] and oxygen-permeable membranes with a hypoxic chamber [74], have been described by others to more closely simulate physiologically relevant lung tumor conditions. Future studies could also look more closely at the development of microscopic KPA tumors following implantation, rather than focusing on visible macroscopic tumors. Further work is necessary to understand the mechanism for the lack of KPA tumor development *in vivo*, and this model provides a unique opportunity to study the effects of PIK3CA ablation in KRAS-driven LUAD.

## 5. Conclusion

We have determined that loss of PIK3CA from KRAS-mutant lung adenocarcinoma prevents the continued growth and progression of tumors *in vivo*, even when implanted directly into the lungs of immunodeficient mice, while markedly increasing sensitivity *in vitro* to oxidative stress from hydrogen peroxide. This provides a model to better understand the fundamental requirements for tumor growth *in vivo* and thus potentially identify new therapeutic approaches.

## Author Contributions

A.L.B designed and performed experiments, analyzed and interpreted data, and prepared the manuscript. G.C designed and performed experiments and analyzed the data. B.R prepared RNA for bulk sequencing. YP.J performed animal surgeries. WX.Z. provided the initial cell line. R.Z.L supervised the project, helped design experiments, interpreted data, and helped prepare the manuscript. H.B supervised the project, helped design experiments, analyzed and interpreted data, and helped prepare the manuscript.

## Funding

This work was supported in part by the Merit Review Award #1I01BX006313 (HB) from the United States (US) Department of Veterans Affairs, Biomedical Laboratory Research and Development Service, the Lung Precision Oncology Program Award # I50CU000162 (HB) from the US Department of Veterans Affairs, Clinical Sciences Research and Development Service, the LUNGevity VA Research Scholar Award #2021-06 (HB), Stony Brook Medicine Pilot Project Program #Dec-2021 (HB), and the National Cancer Institute award #R21CA274425 (RZL).

## Disclaimer

This manuscript does not represent the views of the US Department of Veterans Affairs or the United States Government.

## Informed Consent Statement

Not applicable.

## Data Availability

The RNA-sequencing raw and processed data has been deposited into the Gene Expression Omnibus (GEO) under accession number GSE306343.

## Acknowledgements

The authors would like to thank Lisa Ballou for preparing the KPA cell line and preliminary characterization that made this work possible.

## Conflicts of Interest

The authors have declared no conflicts of interest.

## Supplementary data

**Supplemental Figure 1.**
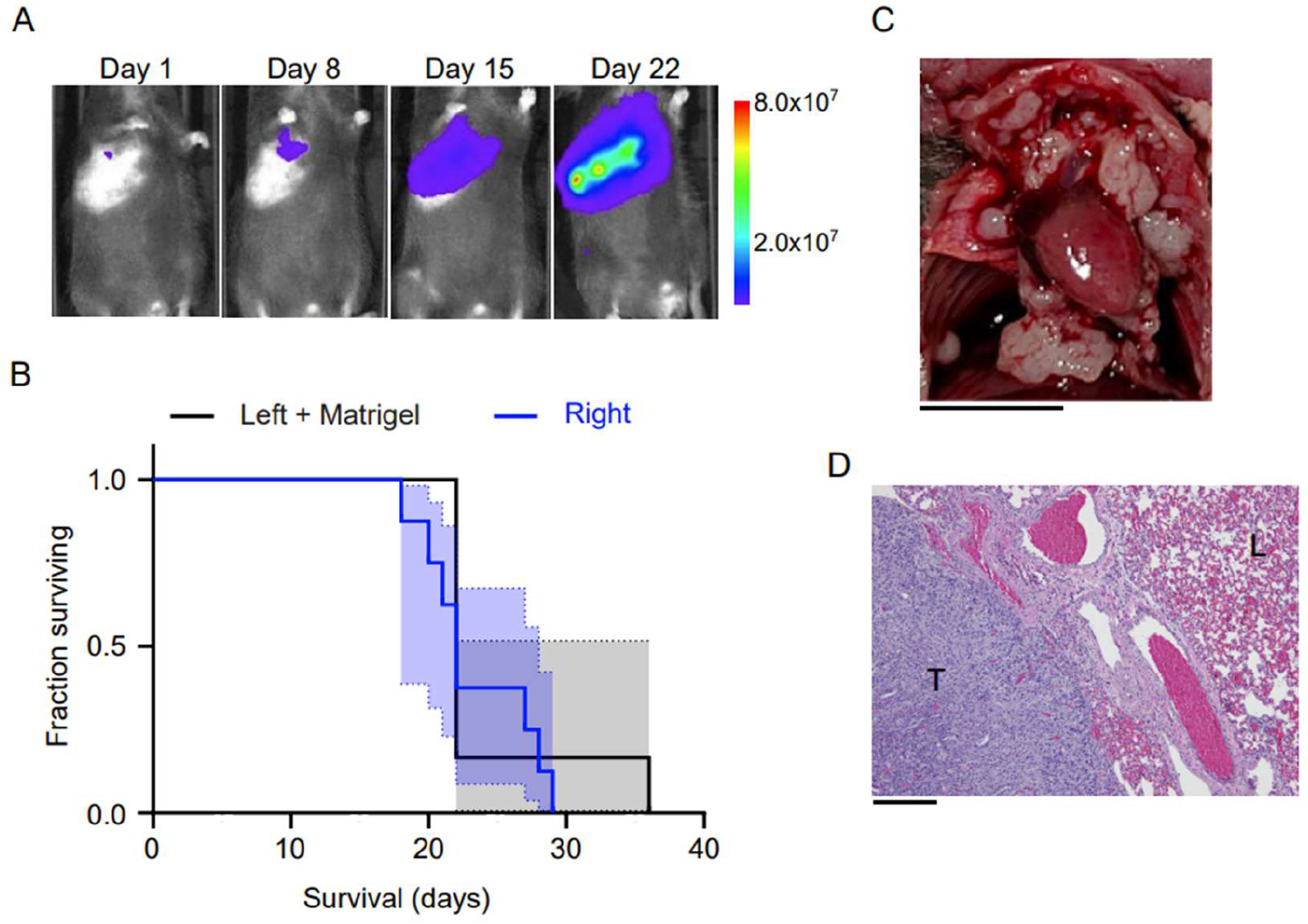
In vivo LUAD model with KP cell line. 5×10^5^ KP cells were implanted into the right (n = 8) or left (n = 6) lung of mice. The cell solution injected into the left lung also contained 500 µg/mL growth factor-reduced Matrigel. (A) Tumor progression was observed with IVIS imaging at weekly intervals points post-implantation. Images from one representative mouse are shown. Scale represents luminescence intensity (arbitrary units). (B) Kaplan-Meier survival curves showing no significant difference and a median survival time of 22 days for both groups. (C) Macroscopic image of chest cavity from one mouse that died of tumor progression. Scale bar represents 10 mm. (D) H&E staining of lung section showing tumor (T) adjacent to normal lung tissue (L). Scale bar represents 200 µm.

**Supplemental Figure 2.**
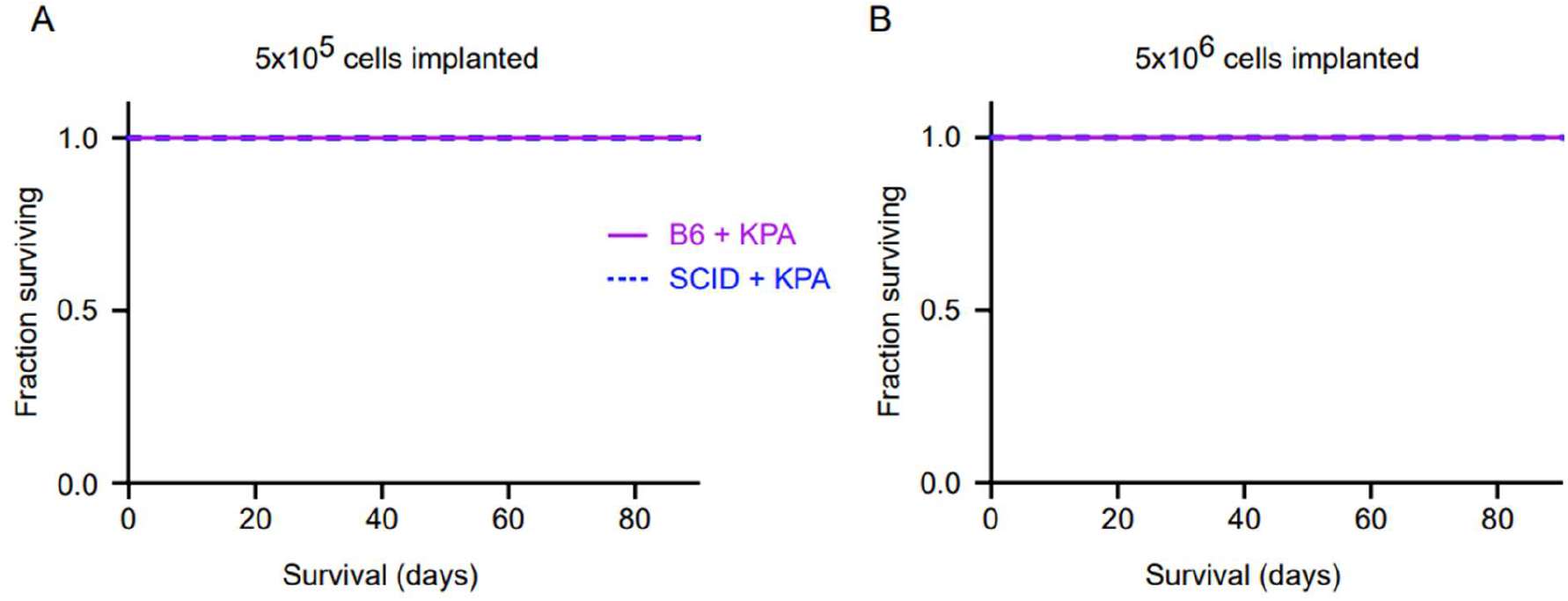
Lack of KPA in vivo tumor progression is not due to adaptive immune response or seeding density. (A) Kaplan-Meier survival curves for B6 or SCID mice implanted with 5×10^5^ KPA cells or KP cells as control in lung (n = 6 per group). Median survival: B6 + KP, 38 days; all mice injected with KPA cells were alive on day 82. p < 0.009, KP versus KPA-injected mice (log-rank test). (B) To test whether simply increasing the number of initial KPA cells may overcome growth limitations, B6 (n = 6) and SCID mice (n = 5) were implanted with ten-fold more (5×10^6^) KPA cells. All B6 and SCID mice were alive on day 82 suggesting growth limitations cannot be overcome simply by increased cell numbers.

## Notes

https://www.ncbi.nlm.nih.gov/geo/query/acc.cgi?acc=GSE306343

### Competing Interest Statement

The authors have declared no competing interest.

